# Structure, identification and characterization of the RibD-enolase complex in *Francisella*

**DOI:** 10.1101/2025.03.02.641097

**Authors:** Xiaoyu Liu, Daniel L. Clemens, Bai-Yu Lee, Roman Aguirre, Marcus A. Horwitz, Z. Hong Zhou

## Abstract

*Francisella tularensis* is a highly infectious bacterium, a Tier 1-select agent, and the causative agent of tularemia, a potentially fatal zoonotic disease. In this study originally aiming to identify anti-tularemia drug targets, we serendipitously determined the atomic structures and identified their components of the native RibD-enolase protein complex in *Francisella novicida*; and subsequently systematically characterized the catalytic functions of the RibD-enolase complex. Originally discovered as individually protein in *Escherichia coli* and yeast, respectively, RibD and enolase are two essential enzymes involved in distinct metabolic pathways, both of which could serve as potential therapeutic targets for tularemia treatment and prevention. Our biochemical validation using pull-down assays confirmed the formation of this complex *in vivo*, revealing that all eluted RibD is bound to enolase, while the majority of enolase remained uncomplexed. Structural analysis reveals unique features of the *Francisella* complex, including key RibD-enolase interactions that mediate complex assembly and β-strand swapping between RibD subunits. Furthermore, molecular dynamics simulations of the ligand-bound RibD-enolase complex highlight localized conformational changes within the substrate-binding sites and suggest a gating mechanism between RibD’s substrate and cofactor-binding sites to ensure efficient uptake and turnover. Despite the physical association between RibD and enolase, enzymatic assays indicated their catalytic activities are independent of each other, thus the complex may have alternative functional roles that warrant further exploration. Our study provides the first structural and biochemical characterization of the RibD-enolase complex, establishing a foundation for further investigations into its functional significance in *Francisella* and potential antibacterial development.

**Significance:** *Francisella tularensis,* a dangerous bacterium classified as a high-priority bioterrorism threat, causes tularemia, a severe and sometimes deadly disease spread from animals to humans. In a study originally aimed at finding new treatments, we accidentally uncovered a unique partnership between two enzymatic proteins—RibD and enolase—in *Francisella*. These proteins, previously known to work alone in other non-pathologic organisms, were found to form a heterodimer of RibD and enolase dimers in *Francisella*. Biochemical experiments confirmed that enolase works both alone and together with RibD in living cells. In addition to the novel heterodimer-of-homodimer configuration, the RibD subunits feature swapped β strands for dimerization as compared to *E. coli* RibD homodimer. The results offer clues for designing future antibiotics against tularemia and beyond.

## Introduction

*Francisella tularensis* is the highly infectious causative agent of tularemia, a potentially fatal zoonotic disease. As a vector-borne disease, insects such as flies, ticks, and mosquitoes are commonly implicated in its transmission between hosts. Human cases can occur through direct contact with an infected animal, bites from arthropod vectors, or ingestion or inhalation of contaminated materials (1). A tularemia vaccine study demonstrated that intracutaneous injection of as few as 10 organisms (2) or inhalation of only 25 organisms (3) of this intracellular gram-negative bacterium can lead to infection. This remarkably high infectivity of *F. tularensis* has led to its classification as a Tier 1 biological agent by the Centers for Diseases Control and Prevention (CDC), due to concerns about its potential use in bioterrorism or biowarfare (4). While *Francisella* infections can be treated with fluoroquinolones and aminoglycosides, other gram-negative bacteria have developed resistance to these antibiotics and there is concern that *Francisella* could acquire such resistance either naturally or by deliberate genetic engineering (5–9), thus posing significant challenges to current therapeutic strategies and emphasizing the importance of development of new strategies for prevention and treatment of tularemia. One promising approach is protein-targeted therapy, which aims to disrupt essential bacterial processes with high specificity. This study explores the discovery, structure and functions of an unexpected *Francisella* complex containing two proteins that play pivotal roles in metabolic pathways and offers insights into their potential as targets for treatment or prevention of tularemia.

The first protein is called RibD (riboflavin biosynthesis protein D), whose homolog was first identified and characterized in *Escherichia coli* during studies on riboflavin (more commonly known as vitamin B₂) biosynthesis (10). Riboflavin is produced through a biosynthetic pathway and serves as the precursor for the biosynthesis of flavin mononucleotide (FMN) and flavin adenine dinucleotide (FAD) (11). These two active forms of riboflavin play essential roles in biological redox reactions crucial for bacterial survival. While riboflavin is biosynthesized *de novo* in microorganisms and plants, the pathway is absent in animals, including humans. This makes riboflavin biosynthetic proteins a potential target for antibacterial drugs (12) as blocking them would not have toxic effects. RibD is a key enzyme in this pathway, responsible for two sequential reactions: the deamination of 2,5-diamino-6-ribosyl-amino-4(3H) pyrimidinedione 5′- phosphate (DARPP) and the subsequent reduction of 5-amino-6-ribosylamino-2,4(1H,3H)-pyrimidinedione 5′-phosphate (ARPP) (12). These reactions are critical for the production of riboflavin, linking RibD to bacterial survival and proliferation. In addition, the product of RibD, 5-amino-6-D-ribitylaminouracil (5-A-RU), is non-enzymatically converted to 5-(2-oxopropylideneamino)-6-D-ribitylaminouracil (5-OP-RU), a microbial metabolite that potently activates mammalian mucosal-associated invariant T (MAIT) lymphocytes, which are capable of killing bacteria-infected cells (13).

The second protein in the newly discovered two-protein *Francisella* complex is enolase. First noted by Otto Warburg in the 1930s in yeast, enolase is a key enzyme in glycolysis, a cytosolic pathway that converts glucose to pyruvate while generating ATP and reducing NAD+ to NADH. In the forward, catabolic direction, enolase catalyzes the conversion of 2-phospho-D-glycerate (PGA) to phosphoenolpyruvate (PEP); while in the reverse, anabolic direction, it converts PEP to PGA during gluconeogenesis (14). Like RibD, enolase is a dimeric enzyme composed of two identical subunits that interact closely to form a functional unit. In addition to its metabolic activity, enolase plays important roles in various nonmetabolic functions (15). It has been observed on cell surfaces as a plasminogen receptor, in the nucleus where it regulates the transcription factor c-Myc, and in the mitochondria where it facilitates tRNA transport for mitochondrial membrane stability (16). Several studies have shown that viral proteins, including those from hepatitis B and C, can hijack enolase to influence viral replication (16). Furthermore, enolase is a component of the RNA degradosome, though only a small fraction (5-10%) of cellular enolase binds to it (17). While its precise function in this complex remains unclear, studies suggest that enolase is not essential for RNA degradation *in vitro* (18). Instead, it may serve a structural or allosteric role, influencing the activity of other degradosome components (19). Due to the abundance of enolase and the high conservation of its amino acid sequence across species, anti-enolase antibodies have been detected in a wide variety of infectious and autoimmune diseases (20). Additionally, studies of natural and synthetic inhibitors of bacterial enolase suggest potential applications in antibacterial therapies (21–23).

In this study, we have determined the atomic structures of, and identified the chemical compositions and interactions within, the native RibD-enolase protein complex from *Francisella novicida*. During an investigation into the protein content of *Francisella tularensis*, we first noticed and subsequently identified a previously unrecognized complex formed by RibD and enolase, two enzymes involved in distinct metabolic pathways. While RibD and enolase have been previously reported in other species to function in separate metabolic pathways, their interaction as a complex has never been documented before. The structural analysis, functional characterization and molecular dynamics simulations of this complex reveal unique features of the *Francisella* complex, offering new insights into its metabolic organization. The structural data provide a foundation for further investigation into the significance of this complex in the bacteria cell and highlight the complex as a potential target for tularemia treatment. Broadly, this serendipitous discovery of the *Francisella* RibD-enolase complex, akin to the approach of cryoID which determines and identifies structures from crude cellular isolates (24), highlights the power of high-resolution imaging techniques like cryogenic electron microscopy (cryoEM) in revealing unforeseen biological relationships.

## Results

### Structure and identification of a RibD-enolase complex in *Francisella*

This study has been developed based on a serendipitous discovery made during the structure-based antibacterial investigation targeting *Francisella*I’s virulence factor, type six secretion system (T6SS). Protein samples purified from *F. novicida* lysate were applied to cryoEM grids for structural analysis. After 2D classification, a small subset of particles exhibiting C2 symmetry, distinct from the dominant population, emerged. These particle images were selected and used for Topaz training (25) for another round of particle picking. Approximately 5% of the particles from the 2D classification were selected and refined, resulting in a 3.05 Å cryoEM map (Fig. S1). The density was of sufficient quality to resolve side chains (Fig. 1D). To identify the proteins within the cryoEM map, we first ran DeepTracer (26). Using the DeepTracer output, a BLAST analysis against the *F. novicida* protein database in Uniprot identified the proteins as RibD and enolase, with molecular weight of 39.6 kD and 49.5 kD, respectively.

**Fig 1.**
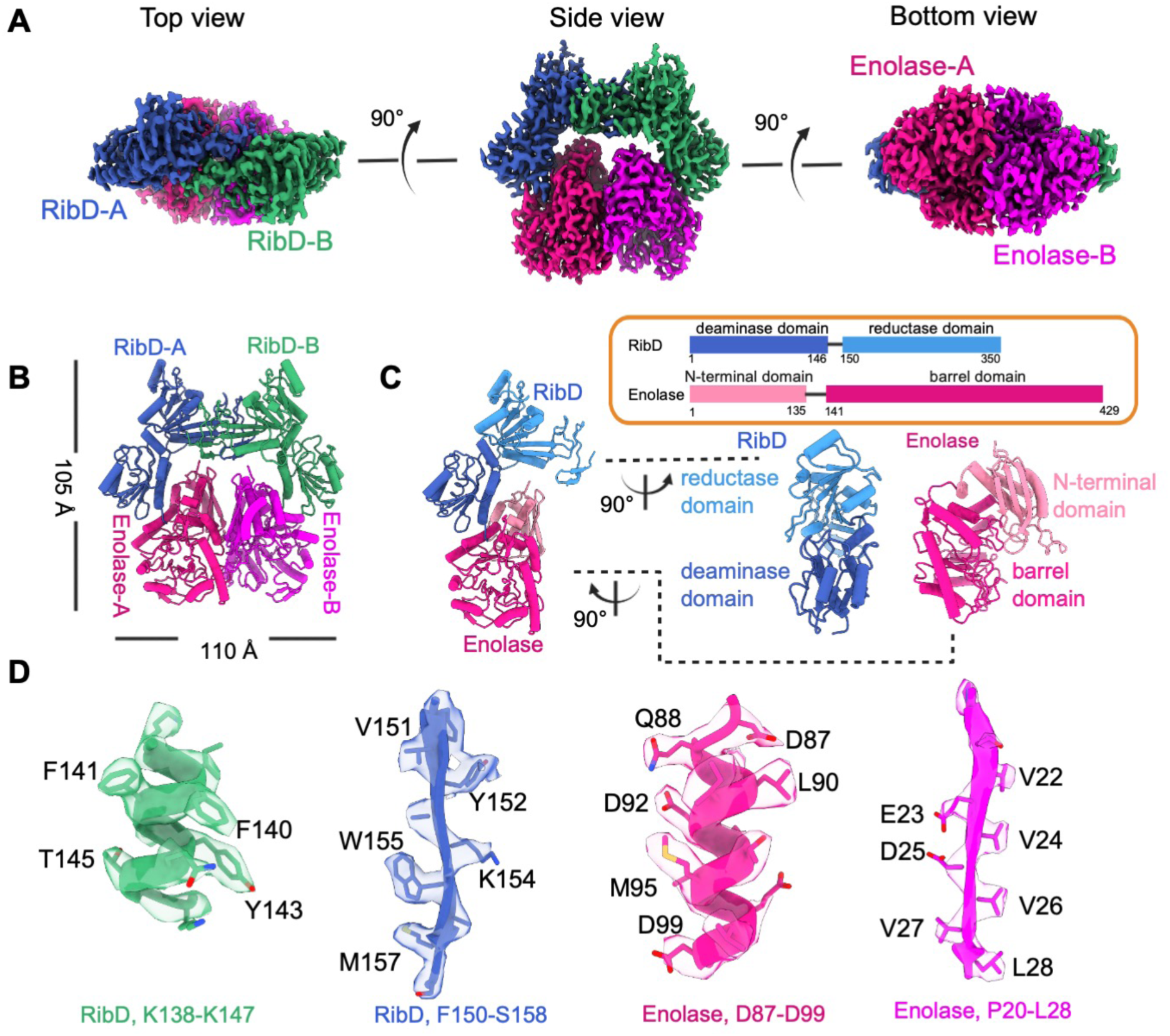
CryoEM structure of RibD-enolase complex. **A.** Three different views of the cryoEM density map of RibD-enolase. **B.** Structure of RibD-enolase complex depicted in cylinder representation. **C.** Domain organization of RibD and enolase. **D.** Local cryoEM density maps.

The structure revealed a heterotetrametric complex consisting of a RibD dimer and an enolase dimer, in a dimer-of-dimer configuration (Fig. 1A, B). The RibD dimer adopts a roof-like conformation, sheltering and interacting with the enolase dimer to form a closed ring. The overall dimensions of the complex are 105 Å × 110 Å × 50 Å. RibD is a bifunctional enzyme with deaminase and reductase activities. It comprises two domains: an N-terminal deaminase domain (1–146) and a C-terminal reductase domain (150–350), connected by a linker region (Fig. 1C). The deaminase domain contains a beta-sheet core formed by five mixed beta-strands, surrounded by alpha-helices. The reductase domain is primarily composed of a beta-sheet with seven parallel strands and a C-terminal β-hairpin, accompanied by several surrounding alpha-helices. Enolase is composed of two domains: a smaller N-terminal domain (1–135) with an antiparallel β-sheet of three strands and four alpha helices and a C-terminal domain (141–429) with a mixed α/β-barrel (Fig. 1C). The deaminase domain of RibD interacts with enolase, forming a key interface within the complex (Fig. 1C). The reductase domain is positioned at the distal end of the RibD-enolase heterodimeric interface.

RibD and enolase are essential enzymes involved in the riboflavin biosynthesis pathway and the glycolysis/gluconeogenesis pathway, respectively. Previous studies have shown both proteins to exist as homodimers (19, 27), but no interactions between them have been reported in any systems. The above structure thus represents the first and direct evidence of RibD and enolase forming a complex in *Francisella*.

### RibD and enolase interact *in vivo*, confirming the physiological relevance of the complex

To verify whether RibD and enolase interact *in vivo*, we performed pull down assays on *F. novicida* cell lysates (Fig. 2A). We prepared *F. novicida* that expresses FLAG-tagged enolase and ALFA-tagged RibD and a control strain that expresses FLAG-tagged enolase without any epitope tag on the RibD. All the proteins were endogenously expressed from their native loci with tagged genes replacing the wild-type genes. Anti-ALFA pull down was performed for lysates of both strains. SDS-PAGE analysis with protein staining revealed two bands corresponding to the molecular weights of enolase and RibD in the strain expressing both proteins with epitope tags. In contrast, no pulldown of enolase was observed in the strain that does not express ALFA-tagged RibD (Fig. 2B). We further confirmed the two bands as enolase and RibD, respectively, by Western blotting utilizing anti-FLAG and anti-ALFA antibodies (Fig. 2C,D). These results demonstrate that RibD and enolase form a complex *in vivo*. In addition, anti-ALFA Western blot showed that the anti-ALFA resin pulls down 100% of the ALFA-tagged RibD but only a small amount of the enolase-FLAG. Thus, in *F. novicida*, RibD exists only in complex with enolase, whereas the majority of enolase is not in complex with RibD. Densitometry of the enolase and RibD bands confirmed that the two proteins are present in a 1:1 ratio, consistent with that revealed in the cryoEM structure.

**Fig 2.**
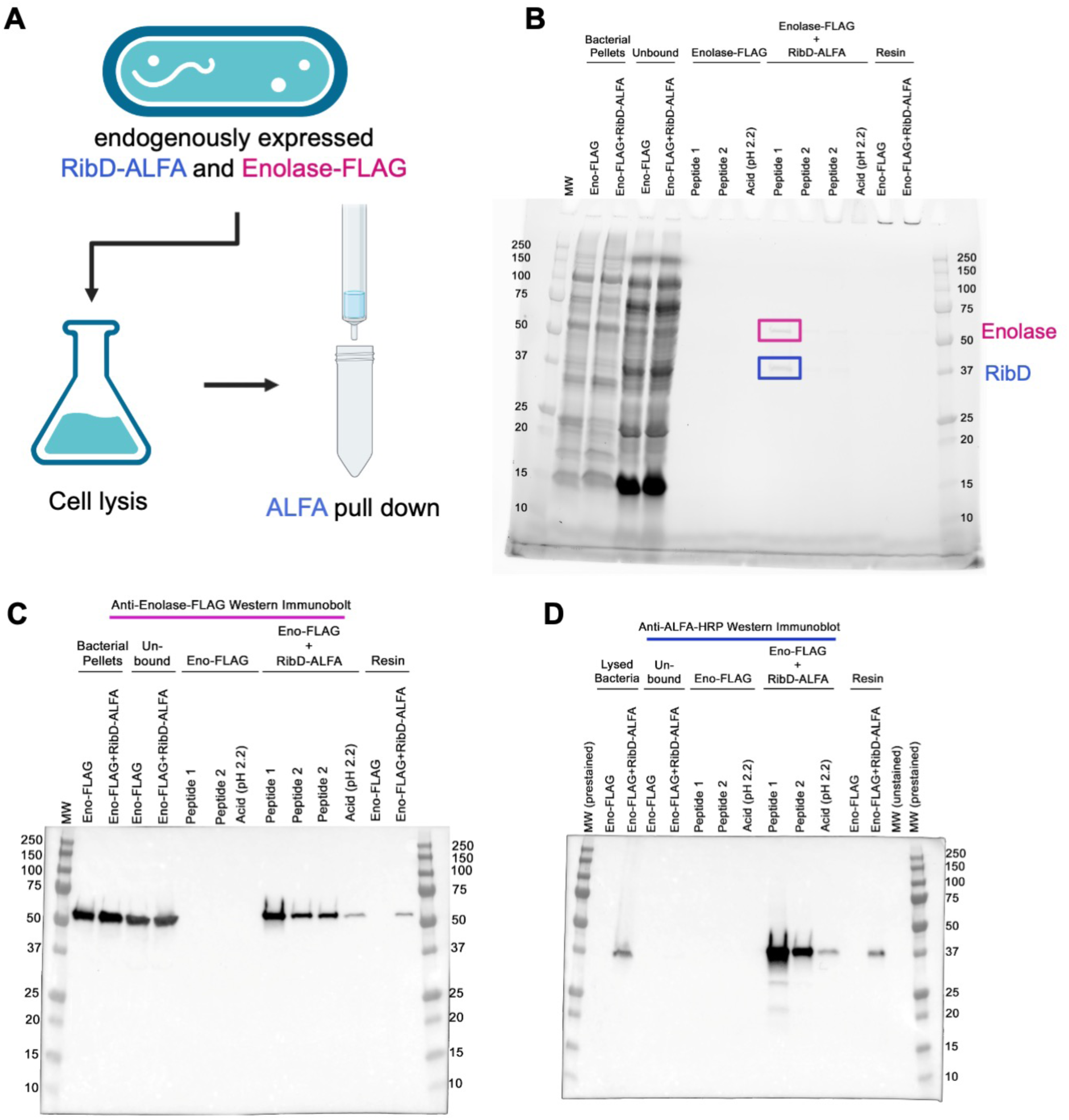
RibD and enolase form a complex in vivo. **A.** Experimental flowchart for pull down assay. **B.** RibD-ALFA and enolase-FLAG are pulled down by Anti-ALFA resin and both are eluted by ALFA peptide. **C.** Anti-FLAG Western Immunoblot confirms that enolase-FLAG is pulled down by anti-ALFA resin in the strain that co-expresses RibD-ALFA and enolase-FLAG. It is not pulled down in the control strain that expresses enolase-FLAG but not RibD-ALFA. **D.** Western Immunoblot is repeated to immunostain for ALFA-tagged RibD.

We further expressed *F. novicida* RibD and enolase as fusions with adenylate cyclase T18 and T25 domains in *E. coli*, and we confirmed the interactions of RibD and enolase by bacterial two-hybrid assay. This assay also demonstrated enolase-enolase and RibD-RibD self-interactions (Fig. S2).

### Interaction interfaces in the RibD-enolase complex

RibD-enolase complex is a heterotetramer consisting of a RibD dimer and an enolase dimer. The two homodimers are brought together through interactions between RibD and enolase subunits. Below, we examine the interactions between the subunits in detail to better understand the structural basis of this assembly.

We first analyzed the interaction interface between RibD and enolase that stabilizes the RibD-enolase complex. The N-terminus of helix α1 from RibD inserts into a hydrophobic groove on enolase, establishing a key interaction. The hydrophobic groove is predominantly formed by residues F350, A354, M136, M353, T384 and C363 of enolase, while the side chains of RibD residues M1, N3, I4, Y7 and Y8 make extensive hydrophobic contacts with the groove (Fig. 3C). The complex assembly is further strengthened by electrostatic interactions between positively and negatively charged residues. Specifically, RibD residues R18 and E42 are in close proximity to enolase residues D87 and R89, forming strong charged contacts (Fig. 3D).

**Fig 3.**
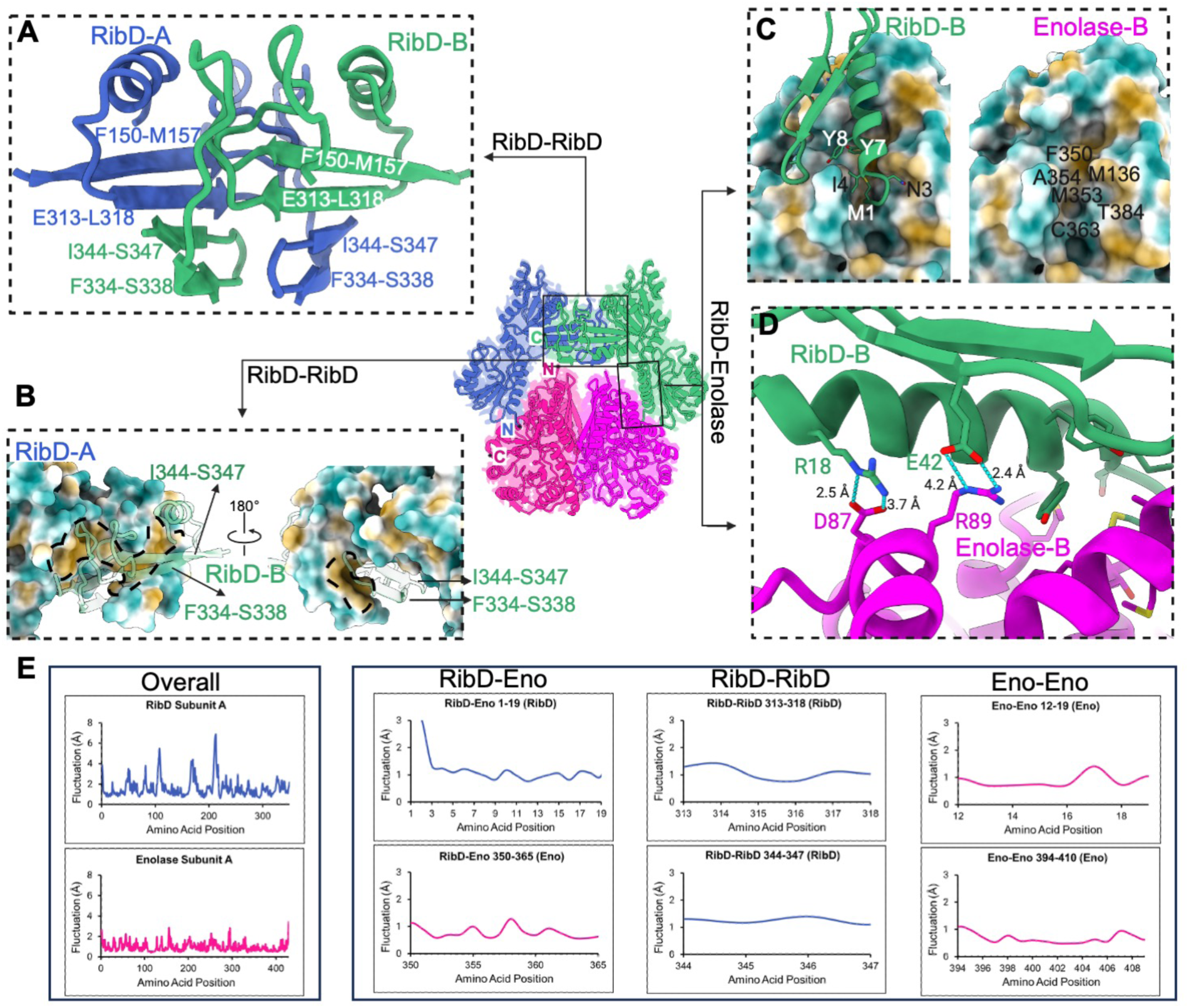
The interfaces between subunits and key interacting residues. **A.** Ribbon representation of interface between RibD-A and RibD-B. **B.** Two views of surface representation with hydrophilic, neutral, and hydrophobic area, indicated in cyan, white, and gold, respectively. The hydrophobic pocket is outlined by dashed line. **C.** Surface representation of interface between RibD and enolase with hydrophilic, neutral, and hydrophobic area, indicated in cyan, white, and gold, respectively. **D.** Charged interactions are shown between RibD and enolase with key residues labeled. **E.** Root-mean square fluctuation (RMSF) calculations of the three interfaces from MD simulations. Low RMSF (≤ 1.5 Å) values are consistently observed with interface-stabilizing residues. Brackets underneath RibD-RibD graph denote the two dimer-swapping regions, 313-318 and 344-347.

The two subunits of the RibD homodimer are held together by their reductase domains involving two interfacing β-sheets (Fig. 3A-B). Key structural elements include β-strands F150-M157 and E313-L318 from one subunit (subunit A) interacting with I344-S347 and F334-S338 from the adjacent subunit (subunit B), forming a four-stranded β-sheet. Similarly, F150-M157 and E313-L318 from subunit B interact with I344-S347 and F334-S338 from subunit A to form another four stranded β-sheet (Fig. 3A). These two swapped β-sheets constitute the primary RibD-RibD interaction interface. The dimeric assembly is mainly stabilized by the extensive hydrogen-bonding network between the β-strand backbones. Additional reinforcements are provided by hydrophobic interactions. Specifically, the strands I344-S347 and F334-S338 from subunit B are located within the hydrophobic pocket formed by subunit A (Fig. 3B).

The enolase dimer interface has been extensively studied across various species, including *E. coli,* yeast and lobster, revealing a conserved structural architecture (19). This conservation is characterized by comparable buried surface areas, surface contact shapes, and charge distributions (19). Notably, enolase dimerization is highly sensitive to ionic strength due to the enrichment of charged residues at the interface. In this study, we show that the *Francisella* enolase dimer exhibits an extensive interaction interface, burying a solvent-inaccessible surface area of ∼3200 Å^2^. Similar to *E. coli* enolase, the *Francisella* dimer interface is enriched with charged residues (Fig. S3B), suggesting a conserved role in dimerization. However, in contrast to *E. coli*, where charged interactions predominate (19), the *Francisella* enolse dimer also incorporates hydrophobic interactions, indicating species-specific variations in dimer stabilization mechanisms.

To investigate the structural stability and dynamic behavior of the RibD-enolase complex in solution, we performed 300-nanosecond (ns) molecular dynamics simulations using GROMACS (28). Structural fluctuations were assessed through Root Mean Square Fluctuation (RMSF) analysis, which quantifies the average deviation of each residue from its mean position throughout the simulation. Overall, RibD exhibited greater fluctuations than enolase (Fig. 3E), suggesting that RibD possesses higher intrinsic flexibility, whereas enolase maintains a more rigid structural profile. Notably, residues involved in complex formation displayed relatively low fluctuations, indicating stable interactions (Fig. 3E). At the RibD-enolase interface, primarily involving RibD residues 1-8 and enolase residues 350-365, the average fluctuation was 1.10 Å. Similarly, the RibD-RibD interface, composed mainly of dimer-swapping residues 313-318 and 344-347, exhibited comparable RMSF values. Enolase demonstrated minimal fluctuations at its homodimeric interface, likely due to its extensive inter-subunit interactions. This contrast in both global and local stability suggests that enolase may stabilize RibD within the complex.

### Comparison of RibD and enolase between *Francisella* and *E. coli* reveals diversification

The above data establish that RibD and enolase form a stable complex in *Francisella*, by contrast, in *E. coli*, RibD and enolase exist independently, with their structures resolved separately (19, 27). To gain insights into the structural differences, we compared the complexed protein structures from *Francisella* with the individually resolved structures from *E. coli*. RibD from *Francisella* and *E. coli* share a majority of their secondary structural elements (Fig. S3). Structural alignment of RibD from *Francisella* and *E. coli* along their symmetry axis revealed a notable difference: the protomers in *Francisella* are positioned further apart than those in *E. coli* (Fig 4A and Movie 1). The wider space between the protomers may facilitate the binding of enolase. Zoomed-in views provide a closer look at the variations observed at the dimerization interface between the two species. In *Francisella*, as described above, the dimerization interface of RibD is predominantly stabilized by swapped β-strands contributed by two protomers (Fig. 4B). In contrast, the dimerization interface in *E. coli* RibD is formed by two β-sheets (27), contributed by subunits A and B, without strand swapping (Fig. 4C). The comparison of enolase structures from *Francisella* and *E. coli* (PDB: 1E9I) (19) reveals a high degree of similarity, with a root mean square deviation (RMSD) of 2.0 Å across all 423 pairs, supporting the understanding that enolases are highly conserved enzymes among bacteria. These structural variations likely reflect functional adaptions specific to the metabolic or regulatory needs of each species.

**Fig 4.**
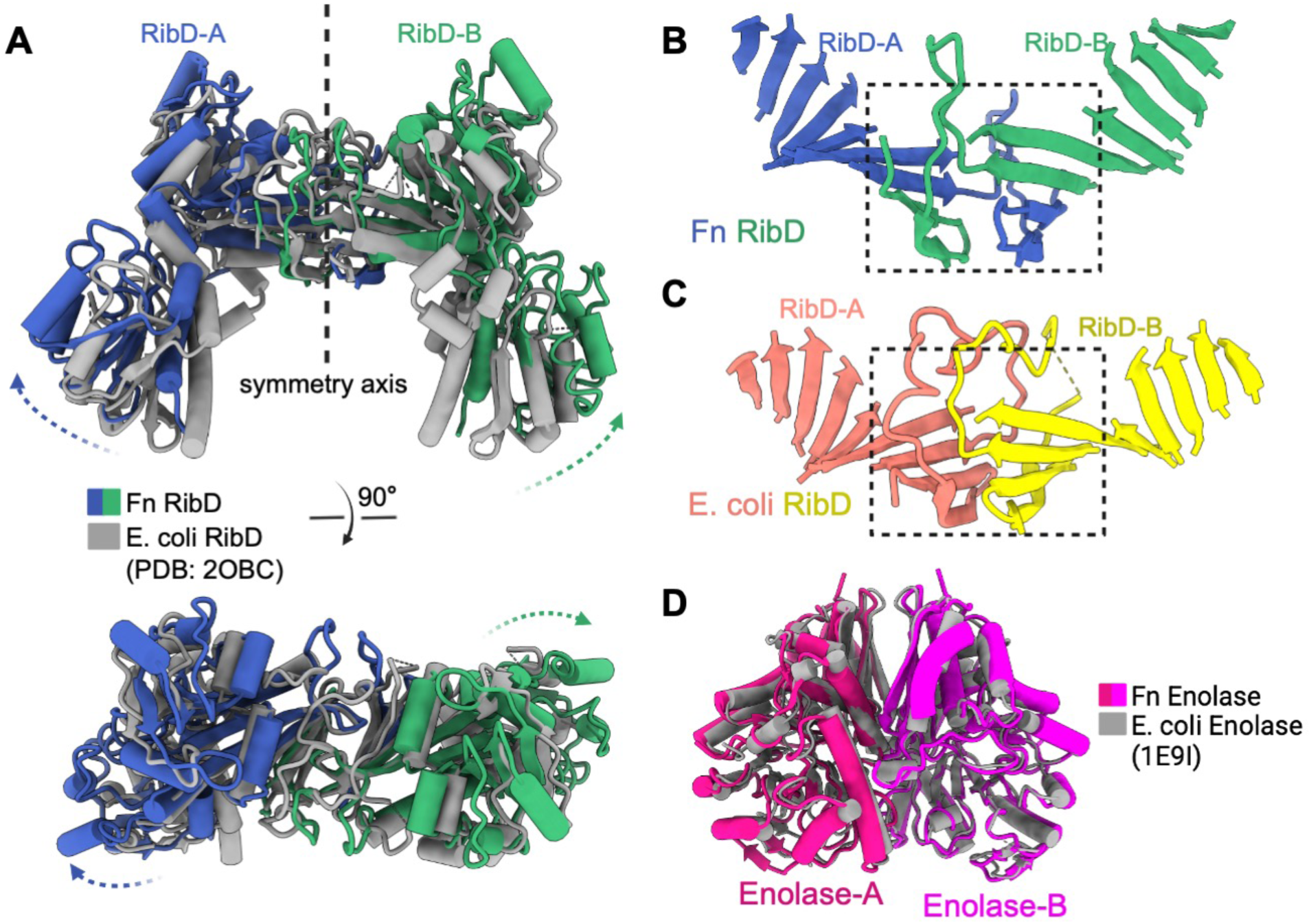
Comparison of RibD and enolase between *Francisella* (Fn) and *E. coli*. **A.** Two views of alignment of Fn RibD and *E. coli* RibD. Fn RibD is colored as blue and green, *E. coli* RibD is colored as gray. The dashed arrows indicate the changes from *E. coli* RibD to Fn RibD. **B.** Ribbon representation of Fn RibD showing the swapped β-strands between RibD-A and RibD-B. **C.** Ribbon representation of *E. coli* RibD showing the interface between RibD-A and RibD-B. **D.** Structure alignment of Fn enolase and *E. coli* enolase showing that the architectures are similar.

### Molecular dynamics simulation show substrate binding and substrate-cofactor occlusion

To gain insight into mechanisms of catalysis, we next carry out computational docking and molecular dynamics simulation. Chai-1 (29) was used to dock DARPP, ARPP, NADPH and Zn^2+^ to RibD subunits and 2-phospho-D-glycerate (PGA or 2PG) and Mg^2+^ to enolase subunits, resulting in 12 total docked ligands (Fig. 5). Afterwards, the experimentally determined structure was overlayed onto Chai-1’s predicted ligands, and the predicted RibD-enolase complex was removed. The protein-ligand complex was then simulated using GROMACS (28) for 300 ns. The RMSF values between the apo and holo simulations were compared to observe any substantial changes between amino acids at their respective binding sites.

**Fig 5.**
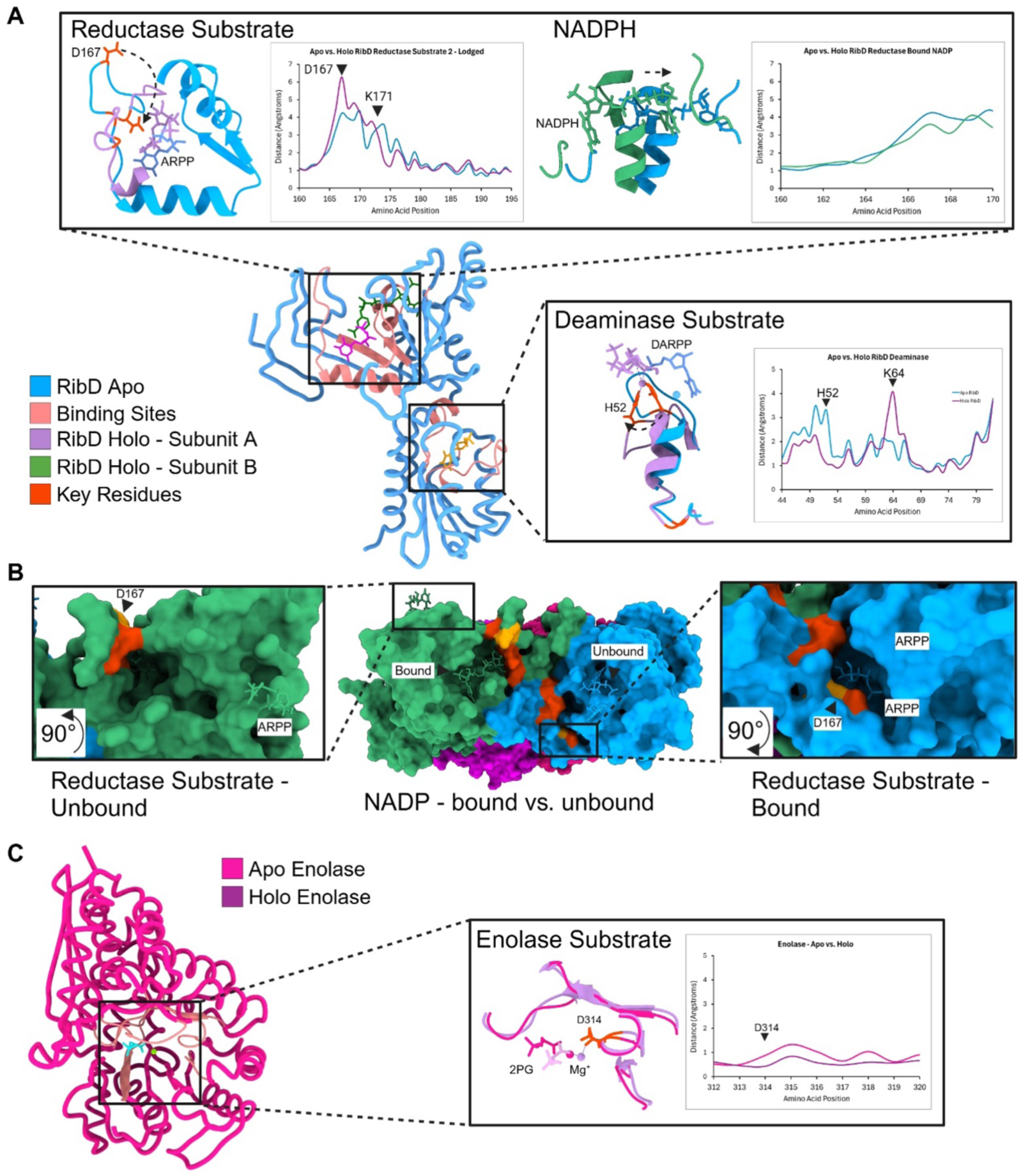
MD simulations RMSF comparisons between apo and holo forms. **A.** Active site fluctuations and local movement to accommodate substrate reductase domain loop 163-174 experiences a large conformational change to bind DARPP, resulting in D167 clamping onto the molecule. However, bound NADPH did not experience large local fluctuations at its binding site. **B**. RibD subunit A (blue) shows a partially dislodged NADP due to the binding site rearrangement from the drastic movement of D167 to the interior of the Reductase substrate binding site to stabilize the substrate (right-hand inset). Conversely, RibD subunit B (green) shows a bound NADP with key residues providing shape complementarity at its active site. This results in the dislodging of the reductase substrate from RibD. **C.** Both enolase subunits bound their substrate in a rigid manner through key residue D314, showing no drastic RMSF changes between its apo and holo forms.

RibD subunit A had DARPP bound throughout the entirety of the 300 ns, but experienced significant local conformational changes occurring at loop 163-174 to better accommodate DARPP within the substrate-binding pocket (Fig. 5A). This loop, containing a crucial D167, clamps down to interact with DARPP (Fig. 5B). On RibD subunit B, NADPH remained bound throughout the entire simulation. There was no noticeable difference between the apo and holo binding sites, with only minor translations from global RibD movements (Fig. 5A). Within the deaminase domain, both substrates remained bound throughout the entire simulation (Fig. S5C), showing large fluctuation differences between the apo and holo forms at residues H52 and K64. While DARPP only undergoes small translational changes, the aforementioned key residues in the deaminase domain have drastic fluctuation changes.

Surprisingly, the simulation trajectories of the holo reductase domain of RibD, containing both DARPP and NADPH, indicated that each subunit only bound one molecule at a time. When NADPH was lodged in one subunit, DARPP would be released, and vice versa (Fig. 5B). This dislodgement is due to the occlusion caused by the highly flexible loop 163-174, since it stabilizes components of both active sites. It cannot accommodate both at the same time, leading to dislodgement of the unattended molecule, but at different time scales (Fig S5A&B). This sheds light into the mechanistic approach for the turnover of fully-realized molecules, allowing for new substrates and reduced NADPH to bind to RibD.

Comparisons between apo and holo simulations of the enolase-2PG binding site do not show any noticeable conformational changes (Fig. 5C). Particularly, holo Asp314, responsible for coordinating with the Mg^2+^ ion, does not exhibit a large mean fluctuation in contrast to its apo form. Both 2PG molecules do not undergo any large conformational changes (Fig. S5D), other than minor translations from their original locations. This relatively low change in both RMSD and RMSF values suggests no substantial conformational differences between enolase’s apo and holo forms. These observations show that enolase lacks flexibility or dynamic capabilities in its substrate-binding pocket, resulting in a small and static active site.

### RibD and enolase in the RibD-Enolase complex catalyze independently

The interaction of *F. novicida* RibD and enolase raised the question as to whether this complex regulates their enzymatic activity. First, we examined whether enolase in complex with RibD exhibits a different enzymatic activity than enolase without RibD. However, we observed that the enolase activity is the same whether or not RibD is present (Fig. 6A). We next purified *E. coli* RibA (Fig. S4A) to examine whether addition of the RibD substrate (generated *in situ* by addition of *E. coli* RibA and GTP) impacts enolase activity, but we found no difference in enolase reaction rate for *F. novicida* enolase vs. the enolase-RibD complex (Fig. 6B).

**Fig. 6.**
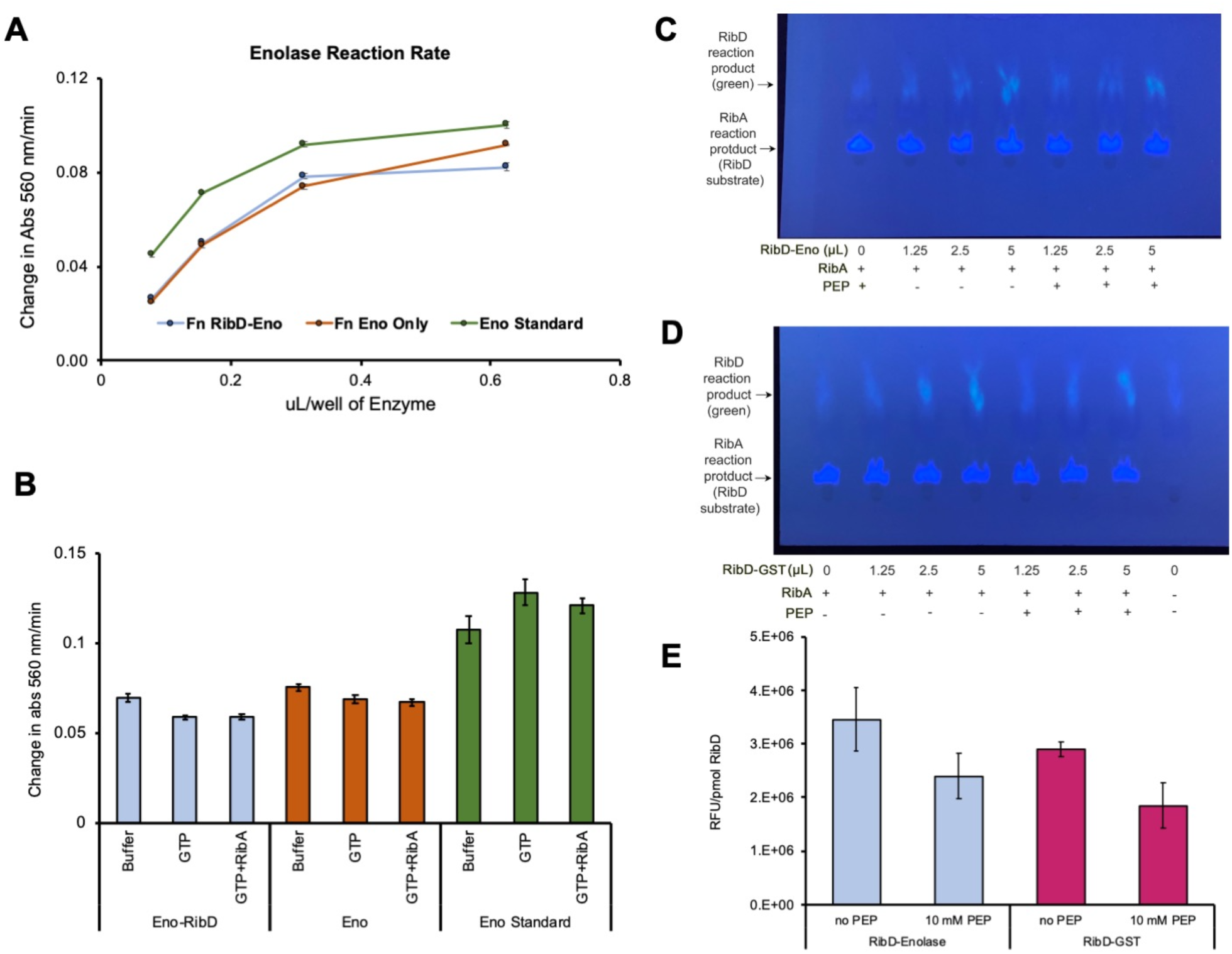
Enzymatic assays show similar activities for individual proteins and the RibD-enolase complex. **A.** Purified *F. novicida* RibD-Eno and enolase without RibD (from stocks of 2.6 and 2.4 µg/mL enolase, respectively) show similar levels of enolase enzymatic activity. **B.** Addition of RibD substrate (generated in situ by addition of GTP and purified *E. coli* RibA) has a minimal effect on enolase enzymatic activity whether or not it is in complex with RibD. Standard enolase (Sigma Millipore MAK178) is included as a positive control in both panels. Data shown are means ± SEM of independent triplicates. **C-D.** Purified RibD-enolase (C) and GST-RibD (without enolase) (D) exhibit similar responses in enzymatic activity to addition of enolase substrate PEP. **E.** Quantification of the RibD reaction product by measurement of fluorescence with excitation at 408 nm and emission at 485 nm. Data indicate means ± standard errors of independent triplicate measurements of relative fluorescent units (RFU) per pmol of RibD.

We next examined whether addition of the enolase substrate, PEP, to the RibD-enolase complex alters the RibD enzymatic reaction rate compared with RibD without enolase. We expressed glutathione S-transferase (GST) epitope-tagged *Francisella* RibD in *E. coli*, purified the recombinant enzyme by affinity chromatography (Fig. S4B), and compared the combined deaminase/reductase enzymatic activities of GST-RibD and RibD-enolase. We found that GST-RibD (without enolase) enzymatic activity is similar to that of RibD-enolase and that the addition of 10 mM PEP has a similar impact on both enzymes (Fig. 6C-E). Thus, we found no evidence that the RibD or enolase regulate each other’s enzymatic activity.

## Discussion

The identification of the RibD-enolase complex was unexpected, underscoring the value of cryoEM in capturing endogenous protein interactions without prior assumptions about their existence. These enzymes have been studied and have been known to function individually in metabolic pathways; therefore, their interaction seen in *Francisella* calls for further studies to determine how the complex form impact the known RibD and enolase catalytic mechanisms, and whether this interaction is specific to *Francisella* or if similar complexes exist in other bacteria.

Molecular dynamics simulations of the RibD-enolase complex with substrates bound suggest a gating mechanism within the broader DARPP/NAPDH binding region, resulting in the occlusion and eventual turnover of both the ligand and cofactor. This dynamic behavior aligns with the previous structural studies of RibD from *E. coli*, which proposed similar conformational transitions (27). The *E. coli* RibD structures captured the loop in the reductase active site in different states: accessible, substrate occluded, and fully occluded with both the cofactor and substrate (27). By integrating our findings with these prior observations, we can corroborate the existence of some of these catalytically viable states, namely the accessible states, in which NADPH and DARPP are able to bind, but then transitions into the dislodging of NADPH (cofactor occluding state), potentially turning over its life cycle due to being oxidized. The other simulated holo RibD subunit keeps the flexible active site loop close to NADPH to stabilize it, akin to the substrate occluded state, but does not allow for proper DARPP binding, resulting in the reductase substrate’s departure from the complex. This simulation result sheds insight into this dynamic region of the RibD substrate, which requires careful modulation of both its substrate and cofactor to ensure efficient uptake and turnover of each.

Formation of the RibD-enolase complex might give rise to new functionalities beyond the RibD and enolase enzymatic activities known for their homodimeric forms. Protein multimerization—the assembly of multiple copies of a single or multiple proteins into larger complexes—serves as an efficient regulatory mechanism in response to changing cellular or environmental demands. For example, multimerization by filament formation, for instance, has been recognized as a key regulatory strategy in the carboxylase superfamily (30); structural studies of filamentous acetyl-CoA carboxylase have revealed two distinct helical filament states: a catalytically active single-stranded helix and an inactive double-stranded helix (31). Multimerization can also diversify enzyme behavior by altering substrate specificity (32), boosting activity (33, 34), or enabling non-catalytic roles like bacterial electron transfer (35), cellular curvature modulation (35), or flagellar motility (36). While we did not observe any evidence that the RibD-enolase complex formation mediates regulatory interaction between the two enzymatic pathways, it is possible that external factors other than the substrates that we examined (e.g., temperature or oxidative stress) might enable such regulation. Given its role as a heat shock protein in yeast (37) and a hypoxic stress response protein in mammalian cells (38), *Francisella* enolase might act as a chaperone to protect and stabilize RibD during stresses encountered in the host cell environment.

## Acknowledgements

We thank Xian Xia for assistance in cryoEM imaging and Kaelyn Y. Feng for editorial assistance. This work was supported by NIH grants (R01AI151055 to MAH and ZHZ, and R01GM071940 to ZHZ). We acknowledge the use of resources at the Electron Imaging Center for NanoSystems supported by US NIH (1S10OD018111) and the US National Science Foundation (DBI-1338135 and DMR-1548924).

## Author contributions

Z.H.Z. and M.A.H. supervised the project. D.L.C. and B.-Y.L. prepared cryoEM sample; X.L. made cryoEM grids, performed cryoEM imaging, and data processing; X.L. built the atomic model and illustrated the structures; D.L.C. and B.-Y.L. prepared bacterial strains, performed biochemistry analysis, enzyme activity assays and bacterial two-hybrid assay. R.A. performed molecular dynamics simulation and analyzed the results. X.L., D.L.C., B.-Y.L., R.A., M.A.H. and Z.H.Z. interpreted the data and wrote the manuscript; and all authors reviewed and approved the paper.

## Methods

### Purification of protein sample

*Francisella novicida* expressing FLAG epitope tagged PdpC was grown in trypticase soy broth with 0.2% cysteine (TSBC), 5% KCl, 0.1 mg/mL FeSO4, and 5 mM betaine to an optical density (540 nm) of 2.0. Bacteria were pelleted by centrifugation (4000 g for 90 min at 4°C), resuspended in 50 mM sodium phosphate, pH 7.4, 1 mM EDTA, 1 mM PMSF, 1 mM NEM, and 1:100 protease inhibitor cocktail (HY-K0010, MedChemExpress), and lysozyme (1 mg/mL) and disrupted by sonication on ice with a probe tip sonicator. The sonicate was clarified by ultracentrifugation (44,400 g for 90 min at 4°C) and NaCl and Mega-9 detergent added to achieve concentrations of 0.15 M and 0.2%, respectively. The supernate was rotated overnight with 0.5 mL of anti-FLAG M2-agarose (Sigma-Aldrich), washed extensively with 50 mM sodium phosphate with 0.15 M NaCl, 1 mM EDTA, and eluted with 3X-FLAG peptide (0.1 mg/mL in the same buffer). The eluate was concentrated with a 100 kDa MWCO filter (Amicon Ultra 15) and used for cryoEM.

### Generation of epitope tagged strains of *F. novicida*

PCR products of the 1.3 kb enolase gene with a C-terminal 3xFLAG tag and 1.2 kb of its downstream genomic sequence, including a repeat of the last 23 nucleotides of the enolase (FTN_0621) gene to preserve the ribosomal binding site, were ligated into pJC84 (39). Similarly, the 1.1 kb *rib*D gene (FTN_0114) with a C-terminal ALFA tag and 0.3 kb of its upstream region and 1.2 kb of its downstream genomic region, including a repeat of the last 24 nucleotides of the *rib*D gene, were cloned into pJC84. Plasmid constructs were confirmed by nucleotide sequencing (Plasmidsaurus) and used for transformation of *F. novicida* U112. Transformants grown on trypticase soy agar containing 0.1% cysteine (TSAC) and 20 μg/ml kanamycin were screened using colony PCR and subsequently counterselected on TSAC containing 7% sucrose. Replacement of the wildtype copy of enolase or *rib*D gene with the epitope tagged gene was validated by Western blotting using FLAG-HRP (Sigma-Aldrich, 1:30000) or ALFA-HRP (Synaptic Systems, 1:4000) antibodies.

### Affinity pull-down of RibD-ALFA

*F. novicida* expressing RibD-ALFA with or without co-expression of enolase-FLAG were grown in 200 mL of TSBC to an optical density (540 nm) of 2, pelleted by centrifugation, resuspended in 50 mM Tris HCl, pH 8, 0.1 M NaCl, 1 mg/mL Lysozyme, 1 mM PMSF, 2 mM NEM, 1:100 protease inhibitor cocktail (MedChemExpress), and 0.2% Tween-20, and sonicated in an ice bath with a probe tip sonicator. The sonicated pellets were clarified by ultracentrifugation (44,400 g for 90 min at 4°C) and the supernates were each incubated with 0.05 mL of ALFA Selector Resin CE (Synaptic Systems) overnight at 4°C with end-over-end rotation, washed with 50 mM Tris HCl, pH 8, 0.1 M NaCl, and eluted in this buffer with 0.2 mM ALFA peptide.

### Affinity pull-down of enolase-FLAG

The clarified lysate of *F. novicida* expressing RibD-ALFA and enolase-FLAG which did not bind to the ALFA-selector resin described above was incubated at 4°C overnight while rotating with 1.5 mL of M2-anti-FLAG agarose (Sigma Aldrich), washed with 50 mM Tris HCl, pH 8, 0.1 M NaCl, and eluted in this buffer with 0.1 mg/mL 3x-FLAG-peptide.

### Cloning and recombinant expression of RibA and RibD

The full-length gene encoding *rib*A was amplified from genomic DNA of *E. coli* NEB5-alpha, digested with *Nde*I and *Xho*I and ligated into expression vector pET28b (Novagen) downstream of a hexahistidine tag and a thrombin cleavage site. The *rib*D gene was amplified from *F. novicida* U112 genomic DNA, digested with *Bam*HI and *Xho*I, and cloned into the vector pGEX-4T-1 for expressing GST-RibD fusion protein with a thrombin cleavage site. The expression plasmids were confirmed by nucleotide sequencing and transformed into *E. coli* BL21(DE3) strain (Novagen). For production of the recombinant proteins, BL21(DE3) cells harboring the pET28a or pGEX-4T-1 construct were grown at 37°C in LB medium until the optical density at 600 nm reached 0.6-0.8 at which time IPTG was added to achieve a final concentration of 0.5 or 1 mM. The cultures were grown at 18°C for 16-20 h and harvested by centrifugation at 3500 g for 20 min and stored at -80°C.

### Purification of *E. coli* histidine-tagged RibA

Bacterial pellets of *E. coli* expressing histidine-tagged RibA were resuspended in 50 mM sodium phosphate, pH 8.0, 0.3 M NaCl, 10 mM imidazole, 1 mM PMSF, and EDTA-free protease inhibitor cocktail (MedChemExpress), sonicated with a probe tip sonicator, and clarified by ultracentrifugation. The clarified supernate was rotated overnight at 4°C with 0.5 mL of Ni-NTA-agarose (Pierce), washed extensively with the lysis buffer and lysis buffer containing 30 mM imidazole and eluted with lysis buffer containing 0.3 M imidazole. Fractions containing RibA were desalted with an Excellulose desalting column equilibrated with 0.1 M Tris HCl, pH 8.

### Purification of GST-tagged *F. novicida* RibD

Pellets of *E. coli* expressing GST-RibD grown as described above were thawed; resuspended in PBS (140 mM NaCl, 2.7 mM KCl, 10 mM Na2HPO4, 1.8 mM KH2PO4, pH 7.3) with 1% TX-100, mM DTT, and 1:100 protease inhibitor cocktail (MedChemExpress); sonicated on ice with a probe tip sonicator; clarified by ultracentrifugation; and incubated with 1 mL of Glutathione-Sepharose 4B (Cytiva Life Sciences) overnight at 4°C while rotating end-over-end. The resin was washed sequentially with PBS with 1% TX-100, and 50 mM Tris, 150 mM NaCl, pH 8.0, and eluted with 10 mM reduced glutathione in 50 mM Tris, 150 mM NaCl, pH 8.0.

### Enzyme activity assays

Enolase enzyme activity was measured by a colorimetric (570 nm) assay using the Enolase Activity Assay Kit (MAK178−1KT, Sigma-Aldrich, St. Louis, MO, USA), which follows the conversion of D-2-phosphoglycerate to PEP by measuring the formation of an intermediate that reacts with a peroxidase substrate.

RibD deaminase and reductase activities were measured at 37°C by a modification of published methods (10, 40). The RibD enzyme assay reaction contained 50 mM Tris, 150 mM NaCl, pH 8.0, *E. coli* his-tagged RibA (0.17 mg/mL), 10 mM GTP, 10 mM NADPH, 10 mM MgCl2, 10 mM DTT, 10 mM glucose-6-phosphate and 1 U/mL glucose-6-phosphate dehydrogenase (Catalog G5885, Sigma Aldrich) for NADPH regeneration. In this system, RibD substrate (2,5-diamino-6-ribosylamino-4(3H)-pyrimidone 5’ phosphate) is generated in situ from GTP by *E. coli* RibA. The RibD deaminase and reductase reactions lead to generation of the reaction product 5-amino-6-ribitylamino-2,4(1H,3H)-pyrimidinedione 5’-phosphate. The reaction is stopped by addition of methanolic diacetyl and boiling for 30 min, which derivatizes the substrate to a blue fluorescent product, 6,7-dimethylpterin, and the reaction product to a green fluorescent product, 6,7-dimethyl-8-ribityllumazine 5’-phosphate. Denatured protein is removed by centrifugation at 14,000 g for 30 min, supernates are evaporated to dryness under vacuum, redissolved in water, spotted onto C-18-W Silica TLC plates (Sorbtech, Norcross, GA) and developed in 20 mM sodium acetate, pH 6, 8% methanol. The blue substrate and green reaction product bands were imaged under long wavelength UV light and areas of the TLC plate corresponding to the green bands were scraped, eluted into 20 mM sodium acetate, pH 6, 80% methanol, and fluorescence measured with a Tecan Spectramax fluorimeter with excitation 408 nm and emission 485 nm.

### Bacterial two-hybrid (BACTH) assay

The genes encoding enolase and RibD were amplified from *F. novicida* U112 genomic DNA or from synthetic gene fragments codon optimized for *E. coli* (Twist Bioscience) and cloned into BACTH expression plasmids (Euromedex) fusing with adenylate cyclase T18-fragment (pUT18 or pUT18c, ampicillin-resistant) and T25-fragment (pKT25 or pKNT25, kanamycin-resistant). The expression constructs were transformed into *E. coli* NEB5-alpha F’ I^q^ (NEB) and verified by restriction digestions and nucleotide sequencing. Two compatible plasmids, expressing the T18 and T25 fusion, respectively, were co-transformed into *E. coli* strain BTH101 by electroporation, and the transformants were selected on LB agar containing 50 μg/ml kanamycin and 100 μg/ml carbenicillin. For detection of protein interactions, antibiotic resistant clones were grown in LB medium containing 1 mM IPTG at 37°C for 16 h, and 3 μl of the cultures were spotted onto LB agar containing 1 mM IPTG and 40 μg/ml X-gal. The agar plates were incubated at 30°C for 48h.

### Graphene grids preparation

The single layer graphene on Cu foil was purchased from Graphene Supermarket. Graphene grid fabrication was done similarly as described in previous work (41). Briefly, MMA EL 6 was used to coat the graphene on copper foil with a spin coater at ∼2500 rpm for 1 min. The graphene on the backside of the copper foil was removed by glow-discharge. Then the copper foil of the MMA/graphene/Cu was dissolved using 1 M ammonium persulfate (APS). The remaining film was transferred to deionized (DI) water and held for 20 min. We used Quantifoil 1.2/1.3 holy girds to harvest the MMA/graphene film and baked the grids for 20 min at 130 °C. The grids were then placed in acetone to dissolve the MMA, followed by transferring grids to isopropyl alcohol (IPA) to remove the acetone residue, and baked for another 20 min to dry. A UV/ozone cleaner was used for 10 min to render graphene grids hydrophilic. The graphene grids were used on the same day they were prepared.

### CryoEM sample preparation and image collection

For cryoEM sample preparation, an aliquot of 3 μL of purified protein sample was applied to prepared graphene grids. The grid was blotted with filter paper (Ted Pella) and then flash-frozen in liquid ethane with an FEI Vitrobot Mark IV (Thermo Fisher Scientific). Freezing conditions were optimized by checking the grids with an FEI TF20 cryoEM instrument. The best grids with optimal ice thickness and particle distribution were obtained with the Vitrobot settings of 8 °C temperature, 100% humidity, 30 s waiting time, 8 s blotting time and 8 blotting force. The prepared cryoEM grids were loaded into a Titan Krios electron microscope (Thermo Fisher Scientific) with a Gatan imaging filter (GIF) Quantum LS and a K3 Summit direct electron detector. The microscope was operated at 300 kV with the GIF energy-filtering slit width setting to 20 eV in super-resolution mode. Movies were collected with SerialEM (42) at a pixel size of Å on the sample level with a total dosage of ∼50 e–/Å2/movie. 39,482 movies were collected.

### Single particle cryoEM reconstruction and model building

The single particle analysis workflow is outlined in Supplementary Fig. S1. Motion correction and defocus value determination were performed in Live CryoSPARC (43). The following data processing was performed with CryoSPARC. 17,070,858 particles were picked by Topaz (25), extracted with boxsize of 300 × 300 pixels and binned 2× to 150 × 150 pixels (2.2 Å per pixel) to speed up the data processing procedure. 407,158 particles were selected from 2D classification. After ab initio reconstruction and homo-refinement, the particles were re-extracted with a box size of 300 × 300 pixels (1.1 Å per pixel). Local CTF refinement, non-uniform refinement with C2 symmetry, and 3D classification were then performed. A total of 356,717 particles from two classes of 3D classification were used for non-uniform refinement, resulting in a final reconstruction at 3.05 Å resolution. The resolution of the map was estimated from the gold-standard Fourier shell correlation criterion, FSC = 0.143. Data collection and processing statistics are summarized in Table S1.

The reconstructed cryoEM map was used to generate a 3D model trace by DeepTracer (26). Based on the sequence model provided by DeepTracer, a BLAST analysis against the *Francisella novicida* protein database in Uniprot identified the proteins as RibD and enolase. Model building is started with the model generated by DeepTracer. The atomic model of RibD-enolase complex was built and refined manually in COOT (44) and further refined in Phenix (45). Refinement statistics of the model is summarized in Table S1. Visualization of the atomic model, including figures and movies, was accomplished in UCSF ChimeraX (46).

### Molecular dynamics simulation and ligand docking

Both apo and holo RibD-enolase files were prepared using the CHARMM-GUI webserver solution builder (47) with the AMBER36 force field at 298K and a pH of 7.5 with GROMACS-compatible outputs. Afterwards, the solvated file was imported into GROMACS (28) and was then subject to pressure and temperature equilibrations for 100 picosecond (ps) each. After verification of proper equilibration, the protein complex was then simulated for 300 nanoseconds (ns). All simulations were run locally using an Nvidia RTX 3070 Ti. The trajectory was then made whole and unwrapped, allowing for direct visualization and analysis using the built-in gromacs tools for RMSF and RMSD calculations. XVG files were manipulated using Microsoft Excel.

For the holo simulation, Chai-1 (29) was used to predict the specific binding regions between the ions and substrates within the RibD-enolase complex. Afterwards, the experimentally-determined model was overlaid into the predicted complex:ligand structure using chimera (46). The predicted structure was then removed, resulting in the ligands being docked into the experimentally determined model. Using the ligand reader function in CHARMM-GUI (47), each individual substrate was extracted and input into the CGENFF server, assigning force fields to the molecules. Finally, the mol2 output files were used in tandem with the Eno/RibD pdb files in CHARMM-GUI’s solution builder, generating a simulation-ready folder.

## Data availability

The cryo-EM density map has been deposited in the Electron Microscopy Data Bank under accession codes EMD-xxxxx. The atomic coordinate has been deposited in the Protein Data Bank under accession codes xxxx.

## Supplementary Figures

**Figure S1.**
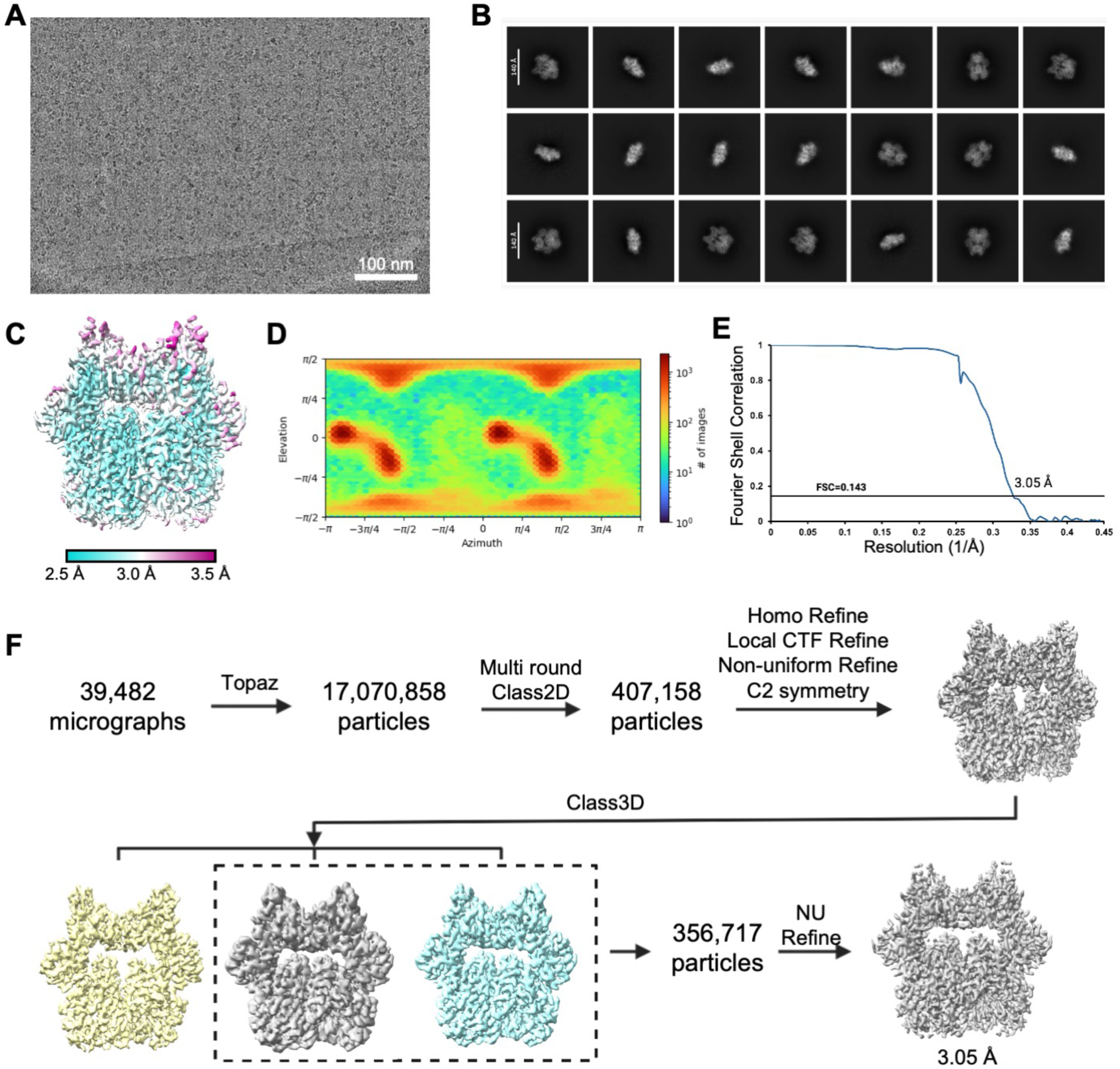
Data processing flowchart. **A.** Motion-corrected cryoEM micrograph. **B.** Representative 2D class averages of RibD-enolase particles. **C.** CryoEM map colored with local resolution. **D.** The angular distribution of the particles for final 3D reconstruction. **E.** Gold-standard Fourier shell correlation (FSC) curve for the cryoEM map. **F.** Flow chart of cryoEM data processing.

**Figure S2.**
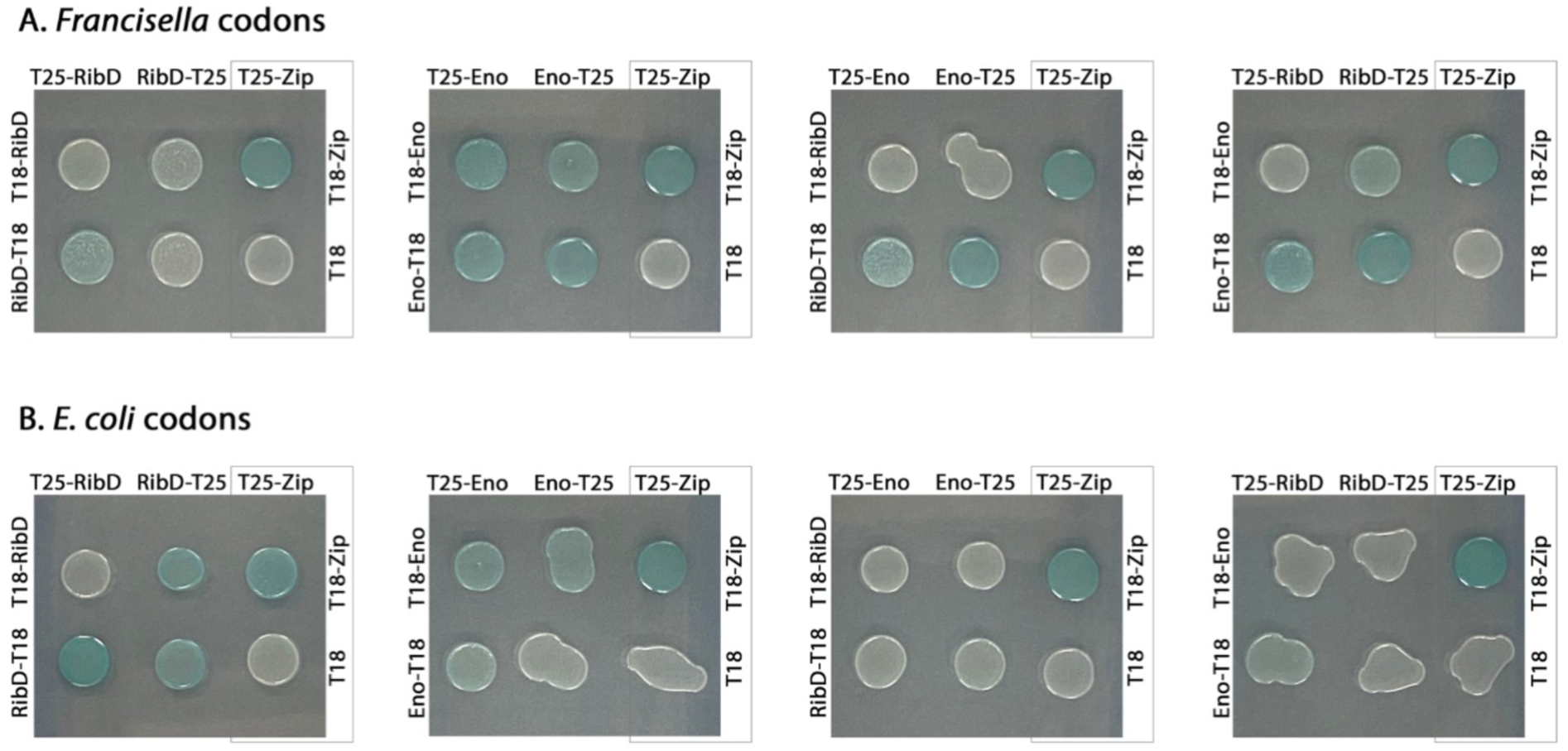
Bacterial two-hybrid assay detects RibD-RibD and enolase-enolase self-interactions and the interactions between RibD and enolase. **A-B.** *F. novicida* RibD and enolase encoded using native codons (A) or *E. coli* codons (B) were expressed as fusions with T18 or T25 catalytic domains of *Bordetella pertussis* adenylate cyclase in *E. coli* BTH101 with IPTG induction. Bacteria were spotted onto LB agar containing X-gal, which acted as a visual indicator to detect protein-protein interactions with blue colonies indicating a positive interaction. Pairwise interactions of RibD and enolase were labeled on the top and left of each spot. Interaction of T18-Zip and T25-Zip served as the positive control, whereas interaction of T18 and T25-Zip served as the negative control.

**Figure S3.**
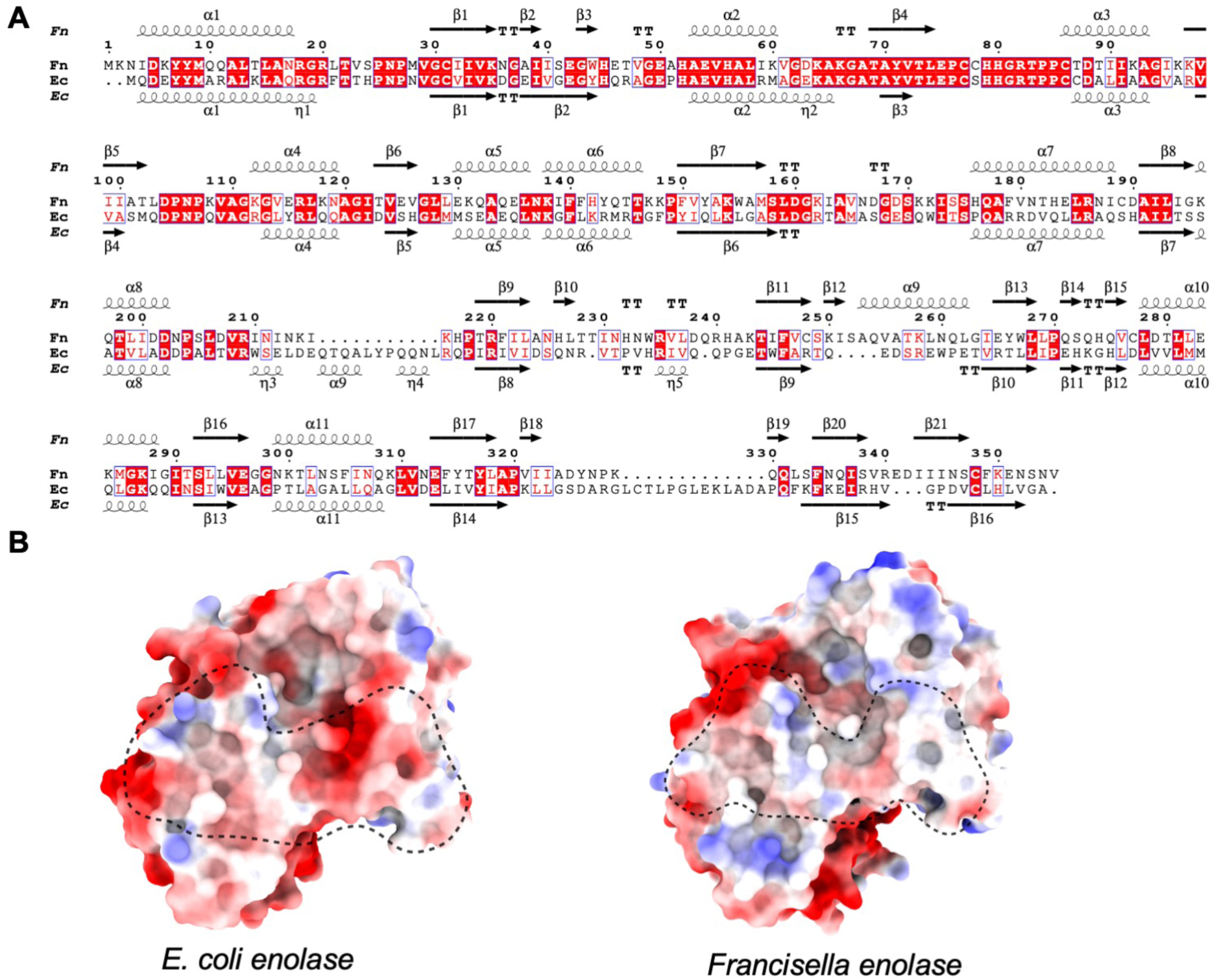
Comparison between *Francisella* and *E. coli*. **A.** Sequence alignment of F. novicida RibD and E. coli RibD. The sequence alignment was done with Clustal Omega and displayed with ESPript 3. Secondary structures of FnRibD determined in this study are shown on top. Secondary structures labeled at the bottom were done according to E. coli structural superposition (PDB accession no. 2OBC). **B.** Electrostatic surface potential of enolase dimer interfaces with positive, neutral, and negative electrostatic potentials indicated in blue, white, and red, respectively. The left panel shows *E. coli* enolase (PDB 1E9I) and the right one shows *Francisella* enolase from this study. The interface area is outlined with a dashed line. The dimer interface is enriched in charged residues.

**Figure S4.**
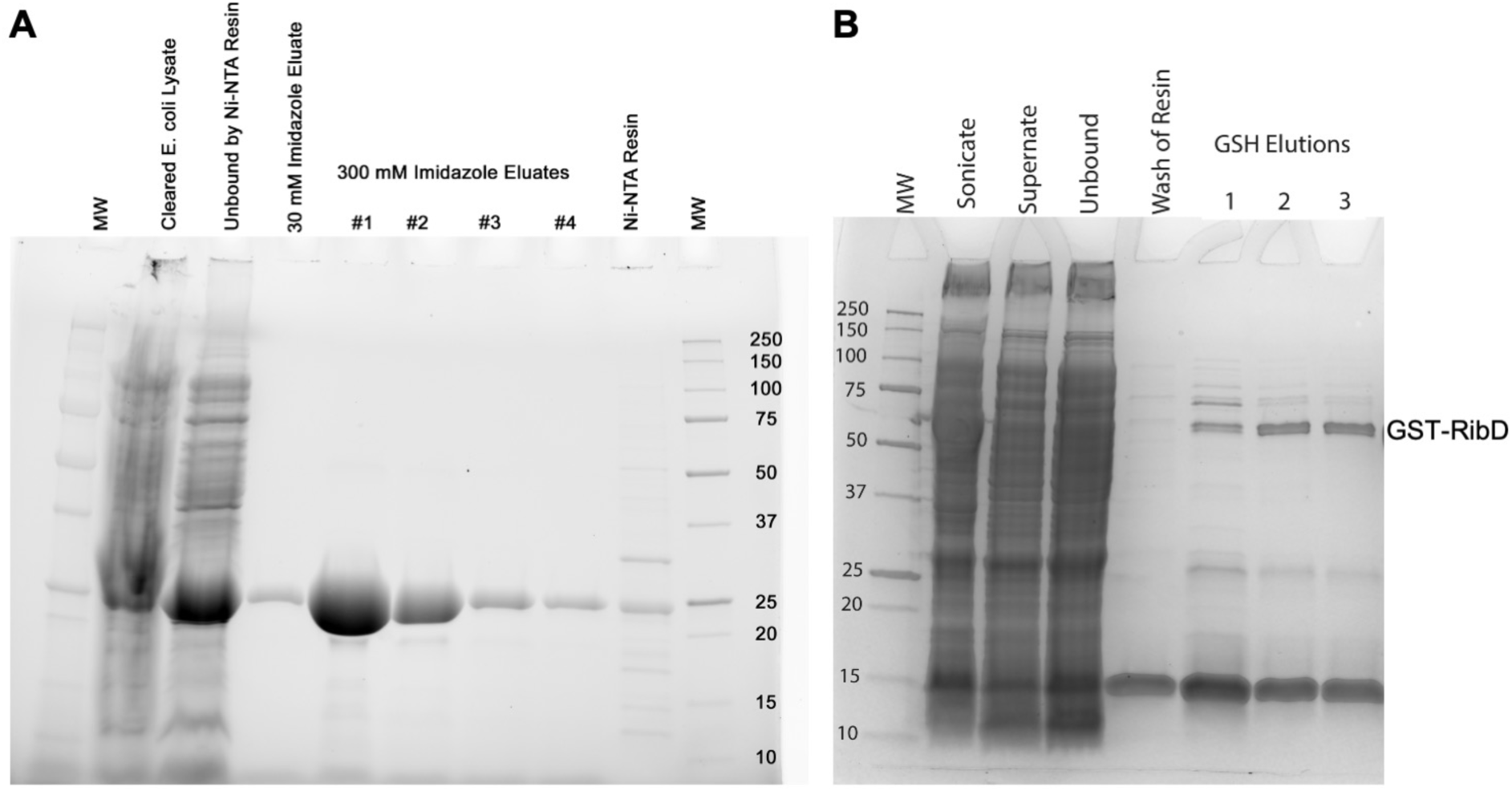
SDS-gel for protein purification. **A.** Purification of *E. coli* histidine-tagged RibA by Nickel-affinity chromatography with evaluation of fractions by SDS-PAGE and stain-free UV imaging. Molecular weight markers are shown on the right in kDa. **B.** Preparation of recombinant GST-tagged *F. novicida* RibD. *E. coli* expressing *F. novicida* GST-tagged RibD was sonicated, clarified by ultracentrifugation, applied to a glutathione-Sepharose column, and eluted with 10 mM GSH. Samples were applied to SDS-PAGE and visualized by staining with Coomassie Blue. The protein band corresponding to the predicted molecular weight of *F. novicida* GST-RibD is indicated. Molecular weight markers are shown on the left in kDa.

**Fig S5.**
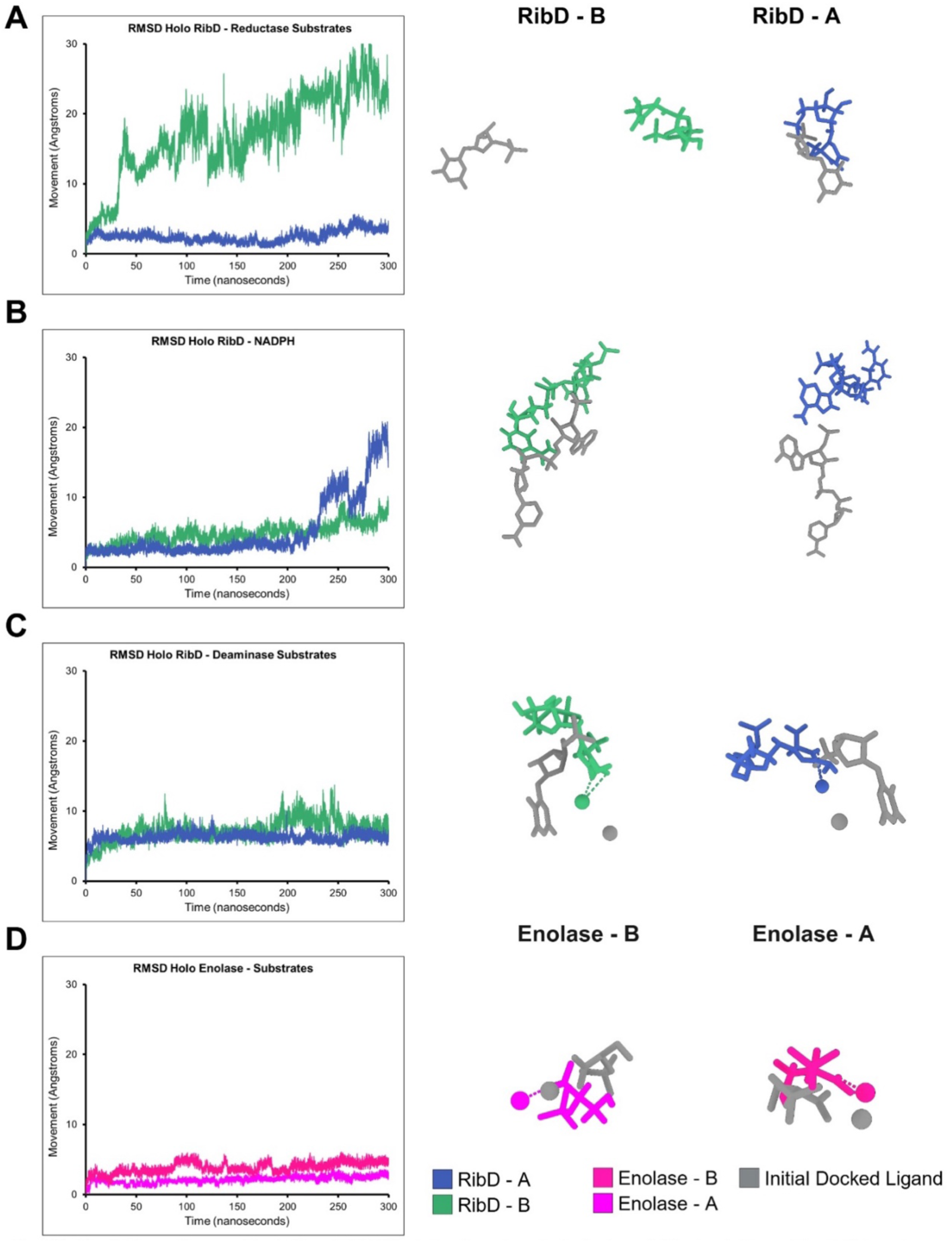
Root Mean Square Deviation (RMSD) plots of each substrate from MD simulations. RMSD plots the deviation of the holo RibD-enolase complex from the superimposed experimental structure throughout the 300 ns. This is shown for **A** The reductase substrates **B** NADPH cofactors **C** Deaminase substrates and **D** enolase substrates. The right hand panels show the distance travel and conformational change of the simulated molecules (color) in comparison to the initial docked molecules (gray).

**Table S1.**
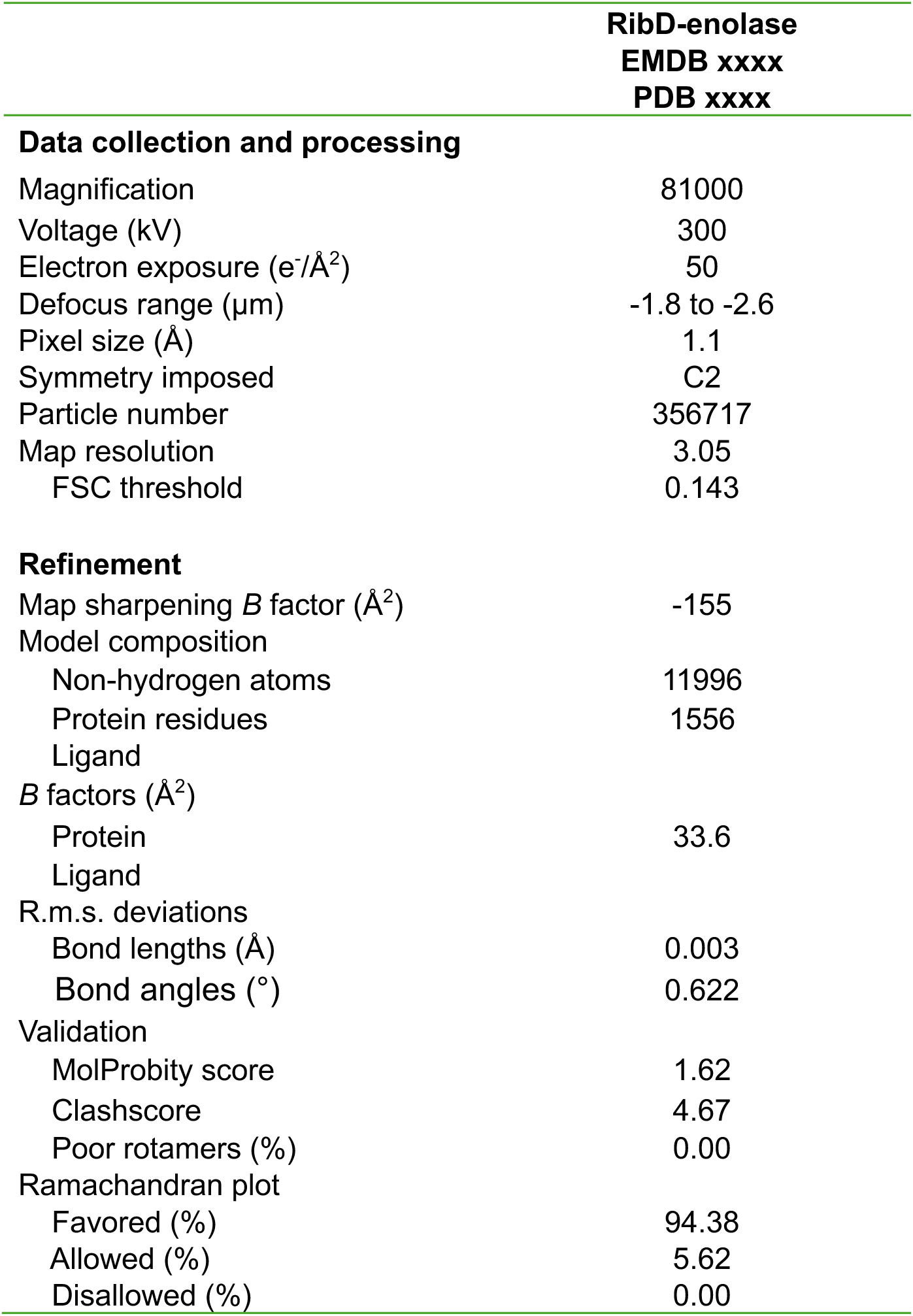
CryoEM data collection, refinement and validation statistics.

|  | **RibD-enolase**<br>**EMDB xxxx**<br>**PDB xxxx** |
| --- | --- |
| **Data collection and processing** |  |
| Magnification | 81000 |
| Voltage (kV) | 300 |
| Electron exposure (e<sup>-</sup>/Å<sup>2</sup>) | 50 |
| Defocus range (μm) | -1.8 to -2.6 |
| Pixel size (Å) | 1.1 |
| Symmetry imposed | C2 |
| Particle number | 356717 |
| Map resolution | 3.05 |
| FSC threshold | 0.143 |
| **Refinement** |  |
| Map sharpening *B* factor (Å<sup>2</sup>) | -155 |
| Model composition |  |
| Non-hydrogen atoms | 11996 |
| Protein residues | 1556 |
| Ligand |  |
| *B* factors (Å<sup>2</sup>) |  |
| Protein | 33.6 |
| Ligand |  |
| R.m.s. deviations |  |
| Bond lengths (Å) | 0.003 |
| Bond angles (°) | 0.622 |
| Validation |  |
| MolProbity score | 1.62 |
| Clashscore | 4.67 |
| Poor rotamers (%) | 0.00 |
| Ramachandran plot |  |
| Favored (%) | 94.38 |
| Allowed (%) | 5.62 |
| Disallowed (%) | 0.00 |

